# The VGCC auxiliary subunit α2δ1 is an extracellular GluA1 interactor and regulates LTP, spatial memory, and seizure susceptibility

**DOI:** 10.1101/2024.12.02.626379

**Authors:** Gerardo Leana-Sandoval, Ananth V. Kolli, Matthew A. Sandoval, Emily Saavedra, Kathy H. Li, Lulu Y. Chen, Alma L. Burlingame, Jorge Ramírez-Franco, Javier Díaz-Alonso

**Affiliations:** Department of Anatomy & Neurobiology, University of California at Irvine, CA, 92617, USA; Center for the Neurobiology of Learning and Memory, University of California at Irvine, CA, USA; Department of Pharmaceutical Chemistry, University of California, San Francisco, San Francisco, CA 94158, USA; Institut de Neurosciences de la Timone, Aix-Marseille Université (AMU) & CNRS, UMR7289, 13005, Marseille, France

**Keywords:** AMPA receptor, cacna2d1, long-term potentiation, voltage-gated calcium channel, object-location memory, hippocampal excitability, transsynaptic organization, AMPAR amino-terminal domain, gabapentinoids

## Abstract

Activity-dependent synaptic accumulation of AMPA receptors (AMPARs) and subsequent long-term synaptic strengthening underlie different forms of learning and memory. The AMPAR subunit GluA1 amino-terminal domain is essential for synaptic docking of AMPAR during LTP, but the precise mechanisms involved are not fully understood. Using unbiased proteomics, we identified the epilepsy and intellectual disability-associated VGCC auxiliary subunit α2δ1 as a candidate extracellular AMPAR slot. Presynaptic α2δ1 deletion in CA3 affects synaptic AMPAR incorporation during long-term potentiation, but not basal synaptic transmission, at CA1 synapses. Consistently, mice lacking α2δ1 in CA3 display a specific impairment in CA1-dependent spatial memory, but not in memory tests involving other cortical regions. Decreased seizure susceptibility in mice lacking α2δ1 in CA3 suggests a regulation of circuit excitability by α2δ1/AMPAR interactions. Our study sheds light on the regulation of activity-dependent AMPAR trafficking, and highlights the synaptic organizing roles of α2δ1.

**Significance statement:** Activity-dependent accumulation of AMPA receptors (AMPARs) at excitatory synapses and subsequent synaptic strengthening underlies long-term potentiation (LTP), forms of learning and memory, and some epilepsies. The “slot model” posits that postsynaptic scaffolding contain “slots” for AMPAR complexes, and that increased synaptic activity augments the availability of slots to accommodate more receptors, thereby strengthening synapses and enabling LTP. The presence of the GluA1 AMPAR subunit amino-terminal domain (ATD) has recently emerged as an additional requirement for LTP. Here we identify the auxiliary voltage-gated calcium channel subunit α2δ1 as a GluA1 ATD interacting protein and provide evidence supporting a role for α2δ1 as an extracellular AMPAR slot regulating activity dependent synaptic AMPAR clustering, excitability, and cognitive function.

## Introduction

AMPA receptors (AMPARs) mediate fast excitatory synaptic transmission in the CNS. Different synaptic plasticity phenomena operate by increasing or decreasing the postsynaptic AMPAR complement, thereby enhancing or weakening synaptic connections, respectively. This includes long-term potentiation (LTP), considered a cellular substrate of learning and memory formation ^1–4^. At hippocampal CA3 Schaffer collateral➔CA1 synapses, as well as other brain synapses, the stabilization of laterally diffusing AMPAR at synapses occurs via interactions between the cytoplasmic domains of AMPAR auxiliary proteins TARPs and PDZ domains in PSD-95 ^5–11^. Super-resolution imaging studies have shown that AMPARs are not uniformly distributed in the PSD. Instead, they are enriched in nanomodules ^12,13^, which number scales with synaptic plasticity^14^. Due to the receptor’s low affinity for its ligand (glutamate), AMPAR activation probability is strongly influenced by the distance from presynaptic release sites ^15–17^. Therefore, alignment of postsynaptic AMPAR nanodomains with glutamate release sites favors receptor activation ^18,19^. How is the transsynaptic AMPAR organization established and maintained? Some synaptic adhesion molecules, including neuroligins, LRRTM 1 and 2, specifically modulate AMPAR transmission, presumably by tuning transsynaptic alignment ^20–23^. A direct contribution of transsynaptic interactions involving AMPARs has been proposed ^15^, but remains to be experimentally established.

Native AMPARs assemble as heterotetramers. Subunit (GluA1-4) composition dictates the receptor’s biophysical properties and trafficking behavior. Specifically, the presence of GluA1 is essential for activity-dependent AMPAR trafficking ^24,25^. Each subunit is comprised of four domains. The transmembrane and the ligand-binding domains are highly conserved among subunits, but the cytoplasmic, carboxy-terminal domain (CTD) and the extracellular, amino-terminal domain (ATD, a.k.a. the N-terminal domain or NTD) are highly sequence-diverse ^26^. For decades, research has focused on the GluA1 CTD, and its activity-dependent posttranslational modifications that modulate synaptic trafficking ^27–29^. More recent research suggests, however, that the GluA1 CTD plays a non-essential, modulatory role in activity-dependent AMPAR trafficking and LTP, at least in CA1 pyramidal neurons (PNs) ^9,30–32^. The ATD is the second most sequence-diverse region. It accounts for roughly half of the protein size, and participates in receptor oligomerization ^33,34^. AMPAR ATDs can induce synaptogenesis ^35,36^ and, as we and others discovered, play subunit-specific roles in synaptic AMPAR trafficking ^37–39^. The emerging model suggests that, at CA1 synapses, the GluA1 ATD is critical for activity-dependent synaptic AMPAR clustering, whereas the GluA2 ATD plays a role in constitutive AMPAR trafficking ^9,37,38,40^.

How does the ATD subunit specifically promote synaptic AMPAR docking? AMPAR ATDs protrude ∼13 nm into the synaptic cleft, approximately half the synaptic cleft width ^26,41^, with GluA1 ATDs typically occupying the outermost positions in the complex ^42,43^. AMPAR ATDs have been speculated to engage in extracellular interactions tuning the receptor’s transsynaptic alignment^16,44–48^. Several AMPAR ATD interactors have been identified ^49^, including pentraxins ^39,50–52^, cadherins ^36^, neuroplastin 65 ^40^, noelins ^53^, and C1ql2 and 3 ^54^. Except cadherins, which can localize presynaptically, AMPAR ATD interactors identified to date are either soluble proteins or localize postsynaptically, and are thus unlikely to exert a direct influence on transsynaptic AMPAR positioning.

To explore extracellular interactors involved in activity-dependent synaptic trafficking, we employed unbiased proteomics. We found that the GPI-anchored voltage-gated calcium channel (VGCC) auxiliary protein α2δ1 is a GluA1 ATD-interacting protein. Our data suggests that presynaptic α2δ1 tunes activity-dependent synaptic AMPAR trafficking, spatial memory, and seizure susceptibility. This study provides evidence of a presynaptic AMPAR slot regulating activity-dependent synaptic AMPAR trafficking, learning and memory, and excitability, suggesting that VGCC auxiliary subunits can act as transsynaptic AMPAR organizing molecules.

## Materials and methods

### Animals

All animal procedures were approved by the Institutional Animal Care and Use Committee at the University of California, Irvine (protocol number AUP-20-156). All animals were maintained in a 12-hour light/dark schedule and had access to food and water ad libitum. C57BL/6-Tg(Grik4-cre)G32-4Stl/J, strain # 006474, ^55^ and B6.129(Cg)-Cacna2d1tm1.1Gfng/J [(α2δ1^f/f^, strain # 030704), ^56^] mice were purchased from Jackson Laboratories and bred to obtain CA3-specific α2δ1 knock out (α2δ1^ΔCA3^) mice. Genotyping was carried out by TransnetYX Inc.

Male and female mice were used indiscriminately in imaging, biochemistry, and electrophysiology experiments; sex was considered as a biological variable in behavior experiments.

### Constructs

pCAGGS-GluA1-IRES-GFP and pCAGGS-GluA2-IRES-GFP were described before ^37^. HA-GluA1 ATD and HA-GluA2 ATD were generated by inserting the HA sequence+stop codon (TACCCATACGATGTTCCAGATTACGCTTAA) after the ATD sequence and cloned into a pCAGGS-IRES-GFP construct. The pCDNA3-α2δ1 construct was purchased from Addgene, where it was kindly deposited by Dr. Lipscombe ^57^, and subcloned in a pCAGGS-IRES-GFP construct. All cloning was done using PCR and In-Fusion HD Cloning System (Clontech).

### Proteomics

AMPAR ATD interacting protein identification was performed with reversed-phase liquid chromatography electrospray tandem mass spectrometry (LC-MS/MS) as described in ^58^. HA-tagged GluA1 or GluA2 ATDs (or mCherry as a control) were transfected using Lipofectamine 2000 (Invitrogen) in 293T cells. 48 hours post-transfection, cells were lysed and the ATDs purified using anti-HA agarose beads (Sigma, #A-2095). Purified ATD-HA beads were incubated with SDS-solubilized P30 mouse brain synaptosome-enriched P2 fractions overnight at 4°C. After washes, pull-downs were eluted in Laemmli buffer (Sigma, #S-3401), separated in SDS-PAGE gels, and silver stained (Thermo Scientific, #24612). After staining, samples were transferred to the UCSF mass spectrometry facility, where bands roughly corresponding to >150 KDa (high MW), 100-150 KDa (med MW) and 50-100 KDa (low MW) were excised from the gel. Proteins in the gel band were reduced with 10 mM dithiothreitol in 25 mM ammonium bicarbonate at 56°C for 1 h, followed by alkylation with 55 mM iodoacetamide in 25 mM ammonium bicarbonate at room temperature for 45 min. in the dark. Samples were then incubated overnight at 37°C with 100 ng trypsin (sequence grade, Promega). The peptides formed from the digestion were further purified by µC18 ZipTips (Millipore) and resuspended in 0.1% formic acid in HPLC water.

The LC–MS/MS analyses were conducted by an LTQ Orbitrap Velos (Velos) mass spectrometer (Thermo Scientific) coupled with a NanoAcquity UPLC system (Waters). During the LC separation, peptides were loaded onto an Easy-Spray PepMap column (75 μm x 15 cm, Thermo Scientific). Following the initial column equilibration in 98% phase A (0.1% formic acid in water) / 2% phase B (0.1% formic acid in acetonitrile) over 20 min., the concentration of phase B was linearly increased from 2 – 30% at a flow rate of 300 nL per minute over 27 min. Then the phase B concentration was increased linearly from 30 – 50% sequentially in the next two min. The column was then re-equilibrated in 98% A / 2% B over 11 min. After a survey scan in the Orbitrap, the top six most intensive precursor ions were fragmented by higher-energy C-trap dissociation (HCD) with a normalized collision energy of 30%. The acquired MS/MS raw data was converted into peaklists using an in-house software PAVA ^59^. The peaklists were then searched against the Uniprot Mus Musculus database (UniProtKB.2017.11.01) using Protein Prospector search engine (http://prospector.ucsf.edu/prospector/mshome.htm).

For each MW range (high, medium, low), we identified specific (i.e., proteins whose NSAF was > 0 after subtracting the Ctrl NSAF value if present in control beads), HA-GluA1 ATD- and HA-GluA2 ATD-interacting proteins. Thus, we compiled a list of specific GluA1 ATD and GluA2 ATD interacting proteins, indicating the total NSAFs (if proteins appear in more than one MW band), as well as the Uniprot predicted subcellular location (Suppl. Fig. 1, Suppl. Tables 1, 2). Afterwards, ATD_GluA1_- and ATD_GluA2_-unique peptides were defined as present in ATD_GluA1_ proteome but absent in ATD_GluA2_ proteome, or vice-versa, as in ^60^.

### Biochemistry

293T cells were cultured in DMEM containing 10% fetal bovine serum, 50 ug/ml penicillin-streptomycin in a humidified CO_2_ incubator at 37°C. Transfections were performed using Lipofectamine 2000 (Invitrogen #11668019,). For Co-immunoprecipitation (co-IP) assays, α2δ1 was expressed together or not with GluA1, GluA2, or HA-GluA1 ATD for 48 h. Cells were then harvested in lysis buffer containing 20 mM Tris pH 7.5, 150 mM NaCl, 1 mM EDTA, 1% Triton X-100, and protease inhibitors (Roche #11836170001). Lysates were incubated at 37°C at 300 rpm for 30 min. and then centrifuged at 10,000 g for 20 min. at 4°C. The lysate supernatants were kept for immunoprecipitation. The supernatants were incubated with either anti-HA beads (Sigma # ROAHAHA) or primary antibody overnight at 4° C followed (in the latter case) by a 1-hour incubation with protein-A agarose beads (Sigma #16266) at 4°C. Beads were then washed three times using washing buffer (20 mM HEPES, 150 mM NaCl, 0.1% Triton X-100, 10% glycerol) at 2,000 g for 1 min. The immunoprecipitates were eluted using 2x Laemmli buffer (Sigma, #S-3401) and boiled for SDS-PAGE and immunoblotting. 1% of the cell lysate was used as input. All samples were assessed by PAGE-SDS electrophoresis. Immuno-Blot PVDF membranes (Bio-Rad, #1620177) were blocked with 5% blotting grade nonfat milk (Lab Scientific, #M0841) in tris-buffered saline buffer with 0.1% tween 20 (Sigma-Aldrich, #P1379). The following primary antibodies were used in co-IP and Western blot experiments: rabbit anti-HA (Cell Signaling, #3724), mouse anti-α2δ1 (Thermo, #MA3-921), rabbit anti-GluA1 CTD (Abcam, #Ab31232), mouse anti-GluA1 CTD (Synaptic Systems, #182 011), guinea pig anti-GluA2 CTD (Synaptic Systems, #182 105), mouse anti-GluA1 ATD (Millipore, #RH95), mouse anti-GluA2 ATD (Millipore, # MAB397). HRP-conjugated secondary antibodies raised against the appropriate species were used: anti-rabbit IgG (Vector laboratories #PI-1000), anti-mouse IgG (Vector laboratories #PI-2000), anti-guinea pig IgG (Millipore Sigma #AP108P). Membranes were incubated with ClarityTM Western ECL (BioRad, #170-5060). Western blots were imaged using BioRad Chemidoc.

### In silico docking analysis

To study the potential interaction between GluA1 ATD and α2δ1, we generated docked models in ClusPro 2.0 server (https://cluspro.bu.edu/login.php?redir=/home.php)^61^ using the full GluA1/GluA2 heterotetramer from pdb:6njl and the full α2δ1 from pdb:7vfv.

Docked models were visually inspected and imposed over the original crystals oriented in pseudo-membranes using OPM server (https://opm.phar.umich.edu/) ^62^ to discard those models not compatible with a trans interaction of GluA1 and α2δ1. Chimera software was used for visualization (https://www.cgl.ucsf.edu/chimera/)^63^. To calculate the binding free energy of the docked models, and to decompose the per-residue contribution, MM/GBSA from the HawkDock server was employed (http://cadd.zju.edu.cn/hawkdock/) ^64,65^. H-bonds were calculated as in ^66^ using https://www.cgl.ucsf.edu/chimerax/docs/user/tools/hbonds.html. For the superimposition of α2δ1 of different species shown in Suppl. Fig. 2, those structures not available as pdb files were generated de novo using AF3 ^67^. The canonical sequence was obtained from Uniprot for the accession numbers O08523 (mouse α2δ1) and P54290 (rat α2δ1). Seeds were automatically generated, using “1338504413” for O05823 and “548886219” for P54290. The first generated model was used for superimposition. json files are available upon reasonable request. For alignment of α2δ1 of different species, “matchmaker” was used in ChimeraX.

### In situ hybridization

15 µm coronal α2δ1^f/f^ and α2δ1^ΔCA3^ mice brain slices were generated following fixation in 4% paraformaldehyde. In situ hybridization (ISH) for α2δ1 was performed using a BaseScope probe (#1331941-C1, Advanced Cell Diagnostics) following manufacturer’s instructions for fresh frozen tissue. Samples were counterstained with hematoxylin (Thomas Scientific) and mounted with Ecomount (Biocare Medical). Samples were imaged with an Olympus CKX53 tissue microscope.

### Confocal microscopy and image analysis

Brains of α2δ1^f/f^ and α2δ1^ΔCA3^ mice were sectioned (40 µm, coronal) following fixation in 4% paraformaldehyde. After blocking tissue with 5% swine serum (Jackson Immuno Research, # 014-000-121) and 2% BSA (Cell Signaling, #9998S) in permeabilizing conditions (0.1% Triton X-100, Sigma-Aldrich, # T8787), samples were incubated overnight at 4° C with the following primary antibodies: GluA1 (rabbit, Cell signaling, #13185, 1:500 dilution), PSD-95 (mouse, Synaptic Systems, #124 011, 1:500), and VGLUT1 (guinea pig, Synaptic Systems, #135 304, 1:500) followed by incubation with Alexa 488 goat anti-mouse (Life Technologies, #A-11001, 1:500), Alexa 647 goat anti-rabbit (Life Technologies, #A21245, 1:500) and Alexa 568 goat anti-guinea pig (Life Technologies, #A11075, 1:500) secondary antibodies for 1.5 hour at RT. Slides were mounted with ProLong Gold Antifade Reagent with DAPI (Cell Signaling Technology, # 8961S).

Confocal images were collected using a Leica Sp8 confocal microscope (Leica Microsystems, Wetzlar, Germany). Images of field CA1 of the dorsal hippocampus were acquired using a 63x oil objective as a series of 20 z-steps, with a z-step size of 1 μm, at a resolution of 1024 x 1024 pixels, scanning frequency of 400 Hz. The optical resolution (voxel size) per image was 180 nm in the xy-plane and 1.03 μm in the z-plane. SIM images were collected using a Zeiss Elyra 7 Lattice-SIM microscope (Carl Zeiss, White Plains, NY). Images of of field CA1 stratum radiatum (SR) of the dorsal hippocampus were acquired using a 63x 1.4 NA Plano-Apo oil objective lens. Images were collected with a z-step interval of 0.273 μm, through a depth of 5.19 μm, covering an area of 64.11 μm x 64.11 μm. Images were acquired at a resolution of 2048 x 2048 pixels. For each focal plane, 13 phase images were collected. The optical resolution (voxel size) per image was 31.3 nm in the xy-plane and 90.9 nm in the z-plane. Images were then processed on the ZEN (black edition) software using ZEN SIM^2^ weak live option.

Analysis of synaptic localization was performed using Imaris 9.9.1 (Bitplane, South Windsor, CT, USA) and MatLab Runtime R2022b (Mathworks, Natick, MA, USA), as previously described ^58^. Briefly, the “Spots” tool was utilized to assign representative three-dimensional ellipsoid shapes to individual synaptic-like puncta of the excitatory presynaptic marker VGLUT1 and postsynaptic marker PSD-95 as well as GluA1. For SIM imaging analysis, the minimum xy and z puncta diameters of GluA1 were set to 0.18 μm and 1 μm, respectively. For confocal imaging analysis, the minimum xy and z puncta diameters of GluA1 were set to 0.5 μm and 0.9 μm, respectively. Then “Background Subtraction” was applied prior to the addition of spots using a Gaussian filtered channel (Gaussian filtered by ¾) minus the intensity of the original channel Gaussian filtered by 8/9th of the punctum radius. A region of interest (ROI) was created to restrict the colocalization quantification to solely within the SR of each image. This ROI spanned an average area of approximately 21,155±1,279 μm^2^ across all samples. ROI was not restricted for the SIM images. The number of spots was adjusted qualitatively using the automatically generated and interactive “Quality” filter histogram to select dense signal while excluding faint puncta likely to be background signal. To ensure an accurate spot segmentation of the underlying puncta determined by size, the “Different Spots Sizes” selection was utilized. Within this setting, the “Local Contrast” tool was used. The corresponding histogram was manually adjusted to ensure each spot covered as much of the puncta as possible. Spots were then rendered. Once optimal settings for each of these parameters were established for the GluA1, VGLUT1, and PSD-95 channels, a batched protocol to automate spot detection on every image was run. For confocal imaging analysis, threshold for colocalization was established at 0.7 μm from the center of neighboring puncta. For analysis of SIM images, threshold for colocalization was established at 0.5 μm from the center of neighboring puncta.

### Electrophysiology

Whole-cell patch-clamp recordings were obtained from CA1 PNs in acute ∼P30 mouse brain slices. 300 µm acute horizontal slices were cut using a vibrating microtome (Precisionary instruments) in ice-cold, oxygenated NMDG recovery solution containing (in mM): 92 NMDG, 2.5 KCl, 1.25 NaH_2_PO_4_, 30 NaHCO_3_, 20 HEPES, 25 glucose, 2 thiourea, 5 Na-ascorbate, 3 Na-pyruvate, 0.5 CaCl_2_·2 H_2_O, and 10 MgSO_4_·7 H_2_O. pH was adjusted to 7.4 and osmolarity to 310-316 mOsm. Slices were then incubated for at least 30 min. at 34°C in artificial cerebrospinal fluid (aCSF) composed of (mM): 119 NaCl, 2.5 KCl, 1 NaH2PO_4_, 26.2 NaHCO_3_, 11 glucose, 2.5 and 1.3 MgSO_4_. aCSF was bubbled with 95% O_2_ and 5% CO_2_ to maintain pH. Osmolarity was adjusted to 307-310 mOsm. For recordings, slices were transferred to a perfusion stage on an Olympus inverted BX51WI microscope and perfused at 2.5 mL/min. with aCSF containing 100 µM picrotoxin. Recording pipettes of 3-6 MΩ resistance were made using borosilicate glass capillaries using a P-1000 puller (Sutter Instruments) and filled with internal solution. The internal solution contained (in mM): 135 CsMeSO_4_, 8 NaCl, 10 HEPES, 0.3 EGTA, 5 QX-314, 4 Mg-ATP, 0.3 Na-GTP, and 0.1 spermine. Osmolarity was adjusted to 290-292 mOsm, and pH at 7.3–7.4. Membrane holding current, input resistance and pipette series resistance were monitored throughout experiments. Data were gathered through a IPA2 amplifier/digitizer (Sutter Instruments), filtered at 5 kHz, and digitized at 10 kHz. Miniature EPSCs (mEPSC) were recorded for at least 4 min. in the presence of picrotoxin (100 μM) and tetrodotoxin (0.5 μM). Series compensation was not performed during data acquisition. Cells with a minimum of 50 mEPSC events were analyzed for event frequency, amplitude, and kinetics. For evoked EPSC recordings, a tungsten bipolar stimulating electrode (Microprobes) was placed in the SR and electric pulses were delivered at 0.2 Hz. AMPAR EPSCs were obtained while holding the cell at −70 mV; NMDAR currents were obtained at +40 mV. The peak evoked AMPAR response and NMDAR component at 100 ms after the stimulation artifact (to avoid contribution of the AMPAR EPSC) were recorded and calculated as the AMPAR/NMDAR ratio. In paired-pulse ratio (PPR) experiments, stimulation was delivered at an inter-stimulus interval of 50 ms. PPR was calculated by dividing the second EPSC by the first.

For long-term potentiation (LTP) experiments, after obtaining a stable baseline, LTP was induced, no more than 6 min. before break-in ^30^, using a theta-burst stimulation (TBS) induction protocol, consisting in four trains of TBS, each train consisting of 5 bursts of spikes (4 pulses at 100 Hz) at 5 Hz applied to the SC fibers at 0.1 Hz, paired with postsynaptic depolarization at 0mV, as in^68^.

Electrophysiology data was gathered and analyzed using Sutterpatch (Sutter Instruments) and Igor Pro (Wavemetrics).

### Behavior

Mice were group housed with littermates. Mice were handled for 1 min for 4 consecutive days prior to all behavioral testing. At the beginning of each testing day, mice were allowed to acclimate to the behavior room for at least 30 min. before the start of the experiment. Behavioral chambers and objects were cleaned and de-odorized between mice. Behavioral scoring was done by a researcher blind to the genotype.

Object Location Memory (OLM) task: Following handling, mice were habituated to a white Plexiglas chamber (30 x 23 x 23x cm) for 5 min. daily for 4 days. On training day, mice were placed in the chamber with two identical objects (2 conical tubes or 2 plastic 3-D printed blocks) and allowed to explore them for 10 min. On test day, 24 hours after the training day, mice were placed in the chamber with either the left or right object displaced to a different location and were allowed to explore the arena for 5 min. Familiar and displaced objects were counterbalanced across mice.

The animal’s behavior was recorded using an overhead camera and exploration time with each object was scored using the criteria described by Vogel-Ciernia & Wood ^69^. Discrimination index (DI) was calculated as follows: (Novel Object Time – Familiar Object Time) / (Novel Object Time + Familiar Object Time) x 100. A DI score of +20 or greater was determined as learning. DI was calculated for both training and test day. Exclusion criteria: Mice that scored ±20 preference for an individual object on training day were excluded from the analysis. Mice that spent less than 3 seconds exploring the objects on training or test day were also excluded. Distance traveled was calculated as: distance traveled (m)/ session duration (min.). Total object exploration was calculated as: total exploration time (sec.)/ session duration (min.).

Novel Objection Recognition (NOR) task: Mice handling and habituation are as described in OLM task. On training day, the mice were placed in the chamber with two identical objects and allowed to explore them for 10 min. The following day (test day), the mice were placed back in the chamber with one familiar and novel object and allowed to explore for 5 min. Discrimination index was calculated as described for OLM.

Forced Alternation Y-maze: The forced alternation task was performed using an opaque Plexiglas Y-maze. Each arm was 36l x 21h x 10w cm. On the training trial, mice were placed into a starting arm, facing the center of the maze, and allowed to explore two of the arms for 5 min., while the third arm was blocked. After an inter-trial interval of 1 min., mice were placed back in the maze at the same starting arm and allowed to explore all three arms for 5 min. The starting arm and blocked arm were counterbalanced across mice. The maze was cleaned and deodorized with 70% ethanol between trials. Total number of arm crossings and time spent in each arm was scored using a mouse tracking software (Any-Maze, Stoelting Co). Mice were required to enter an arm with at least 2/3 of its body to be considered a crossing. DI was calculated as Novel Arm Time / (Novel Arm Time + Non-Starting Arm) x 100 (Wolf et al., 2016).

Forced Swim Test: Mice were individually placed in a clear plastic cylinder (10h x 9d in) filled with water at 23°C for 5 min. The total time spent immobile in the last 3 min. of the task was individually scored by a researcher blind to the genotype. Floating and idle swimming were considered immobility (Can et al., 2012).

Light/Dark Transition Test: The light-dark apparatus consisted of an opaque acrylic box (42 x 21 x 25 cm) divided into two compartments (2/3 light, 1/3 dark) with a small opening connecting the two chambers. The light compartment was made of opaque white walls and lit by an overhead lamp, while the dark compartment was unlit and made of black non-transparent acrylic walls. Mice were first placed in the light compartment allowed to freely explore both chambers for 10 min. The time spent in each chamber, number of crossings, and the latency to enter the dark chamber was recorded using Any-Maze (Stoelting Co).

### PTZ susceptibility test

Pentylenetetrazol (PTZ, Sigma-Aldrich) was dissolved at a concentration of 22.5 mg/kg in 0.9% saline and administered intraperitoneally to ∼3 month-old α2δ1^f/f^ and α2δ1^ΔCA3^, individually housed in plexiglass cages every 10 min., as described in ^70^. Mice were injected with the PTZ-containing solution every 10 min., with consecutive injections of PTZ until generalized seizures occurred. This was considered the end of the experiment. Seizure severity was monitored following a modified Racine scale (Shimada & Yamagata, 2018). Briefly, 0: normal behavior, 1: motionless, laying on stomach, 2: head nodding, 3: stiff tail, forelimb and hind limb myoclonus, 4: rearing with forelimb myoclonus, falling on its side, 5: tonic-clonic seizure, jumping, death. All the procedure was video-recorded and re-analyzed later by an experimenter blind to the experimental groups, to determine the precise onset of generalized seizures. There was no statistically significant difference in weight or sex ratio between the different groups of mice.

### Statistical Analysis

Data analysis throughout the study was done blind to the experimental condition when possible. Results shown represent the mean ± SEM. The number of independent experiments or biological samples, and the statistical test employed, are indicated in every case. Statistical analyses were performed using GraphPad Prism 9 and SutterPatch software.

## Results

### A putative extracellular AMPAR slot

We and others previously found that truncation of the GluA1 ATD blocks synaptic AMPAR trafficking and LTP ^37,38,40^. Based on these results, we hypothesized that the GluA1 ATD contributes to synaptic AMPAR docking by interacting with extracellular elements in the cleft. Previous proteomic screens have identified numerous AMPAR-binding proteins ^71,72^. The vast majority of them are either transmembrane or cytoplasmic. Therefore, they most likely interact with AMPAR in cis, with minimal, if any, contribution of the GluA1 ATD. To identify specific transsynaptic AMPAR interactors, we incubated mouse forebrain homogenates with recombinant, HA-tagged, GluA1 and GluA2 ATDs (Fig. 1A). We then performed an unbiased proteomic screen to pinpoint specific GluA1 ATD - vs GluA2 ATD-interacting proteins. Given its importance in AMPAR trafficking ^37^, we focused on specific and unique ATD_GluA1_-binding proteins with extracellular or transmembrane topology (Fig. 1B, C, Suppl. Fig. 1). Among the candidates found (Suppl. Table 1), we focused on the VGCC auxiliary subunit α2δ1, encoded by the intellectual disability- and epilepsy-associated gene CACNA2D1 ^73,74^ for further analysis, based on i) its extracellular topology, associated with the presynaptic membrane via a GPI anchor ^75^, ii) its selectivity for GluA1 ATD vs. GluA2 ATD (Fig. 1C, Suppl. Table 1), iii) its proximity to glutamate release sites owing to its association with pore-forming VGCC α subunits ^75,76^, and iv) its involvement in transsynaptic signaling ^77,78^. First, we verified the direct interaction using co-IP of recombinant α2δ1 and GluA1 in 293T cells (Fig. 1D). In this assay, α2δ1 can also interact with GluA2, albeit with qualitatively lower affinity (Fig. 1E). We then tested whether the GluA1 ATD alone can bind α2δ1, and found that the interaction is at least as strong as with the full-length GluA1 subunit (Fig. 1F). Finally, we pulled-down GluA1 from mouse forebrain synaptosomal fractions and confirmed that the interaction with α2δ1 occurs in vivo (Fig. 1G).

**Figure 1.**
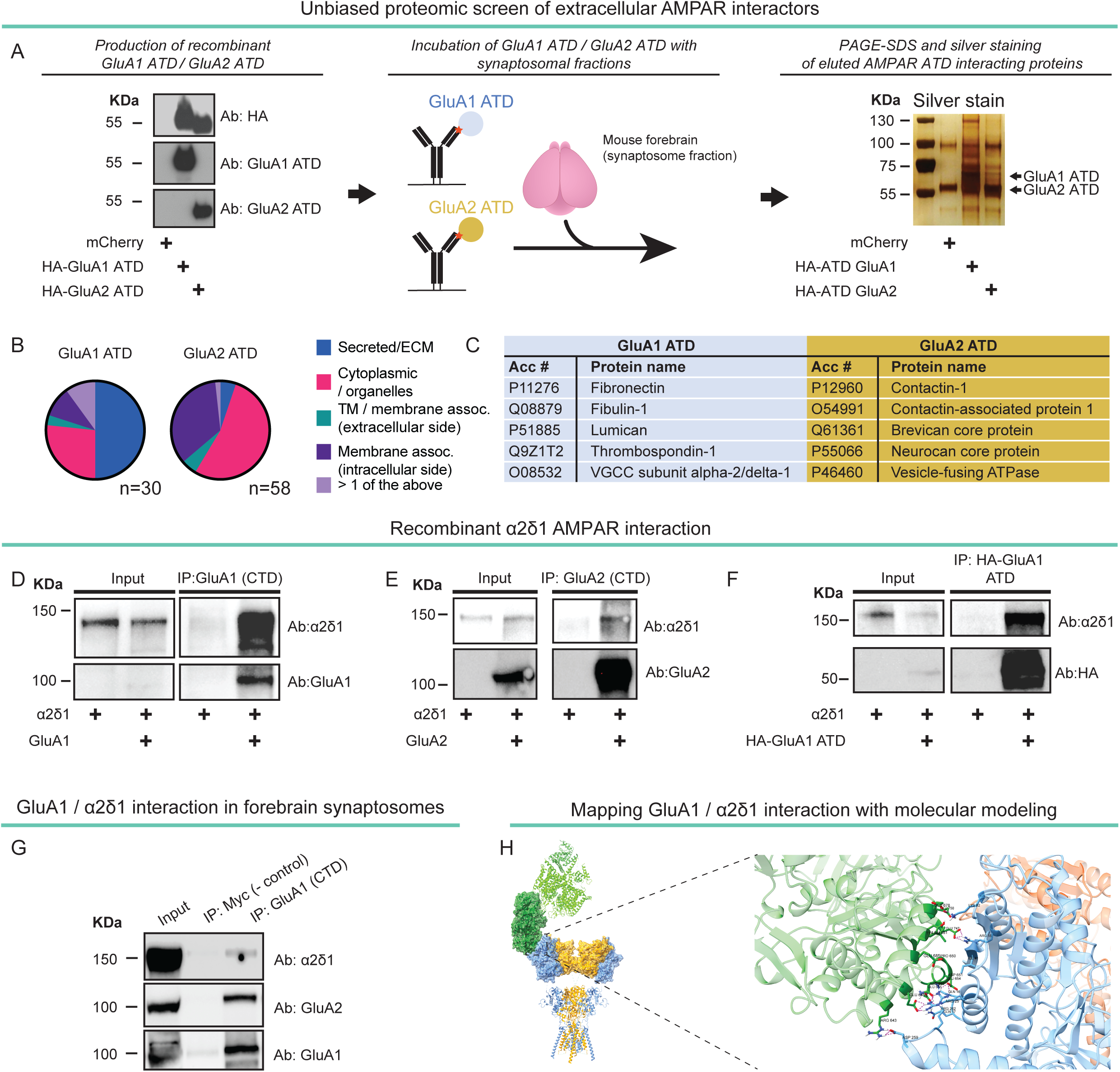
Identification of AMPA receptor ATD interacting proteins. A, Left, purified recombinant HA-GluA1 ATD and HA-GluA2 ATD produced in 293T cells, then used as bait in pull-downs. Center, schematic of pull-down assay after incubation of recombinant ATDs with whole mouse brain lysates. Right, silver stain of proteins eluted after pull-down and PAGE-SDS. B, Subcellular localization of specific and unique GluA1 ATD and GluA2 ATD interactors. C, Partial list of GluA1 ATD (left column, blue) and GluA2 ATD (right column, yellow)-interacting proteins in the mouse brain identified in proteomic screen. D, co-IP analysis of the interaction between recombinant α2δ1 and GluA1 (n≥3). E, co-IP analysis of the interaction between recombinant α2δ1 and GluA2 (n≥3). F, co-IP analysis of the interaction between α2δ1 and HA-GluA1 ATD (n≥3). G, GluA1 / α2δ1 interaction in mouse brain homogenates after pull-down with GluA1 CTD antibody. GluA2 is used as positive control (n=3). H, Left, Molecular model of the ATD of AMPAR (pdb: 6njl) docked to α2δ1 (pdb: 7vfv) in complex with the VGCC allowing a trans interaction obtained in ClusPro docking server. GluA1 subunits are show in light blue, GluA2 subunits are shown in yellow. α2δ1 is shown in dark green, the rest of the VGCC complex in light green. Right, the region boxed in the H is depicted at higher magnification, highlighting some of the residues involved in the interaction between GluA1 ATD and α2δ1. Potential intramolecular H-bonds between the selected residues are indicated in magenta. Rendering of the molecular complexes was performed in ChimeraX.

AMPARs interact with postsynaptic α2δ1 (i.e., in cis) in dorsal root and hypothalamic synapses ^79,80^. However, at most CNS synapses, α2δ1 localizes primarily presynaptically ^81^, and has been shown to act as a transsynaptic organizer at both inhibitory and excitatory synapses ^77,78^. However, the precise interactions that fulfill this role are unknown. To explore the possibility that presynaptic α2δ1 directly interacts with AMPAR in trans, thus potentially contributing to AMPAR localization in the transsynaptic nanocolumn, we generated docked models in ClusPro 2.0 server ^61^ using the full GluA1/GluA2 heterotetramer ATD crystal structure from pdb:6njl ^42^ and the full CaV2.2/α2δ1/β1 complex crystal structure from pdb:7vfv ^82^. Interestingly, some of the docked models obtained were compatible with a transsynaptic interaction between α2δ1 and the GluA1 ATD enabling the alignment of AMPARs and VGCCs (therefore, with putative glutamate release sites, Fig. 1H, Suppl. Fig. 2B). Docked models compatible with an interaction in trans were imposed over the crystals oriented in pseudo-membranes using the OPM server ^62^, and the residues involved in putative contact surfaces between α2δ1 and GluA1 ATD were mapped (Fig. 1H, Suppl. Table 2). We confirmed that the α2δ1 interface interacting with the GluA1 ATD is conserved in other mammalian species for which crystal structures are unavailable, using structures predicted by Alphafold 3 (Suppl. Fig. 2), suggesting that this putative interaction is conserved.

### Presynaptic loss of α2δ1 affects synaptic plasticity, but not basal transmission, at hippocampal Schaffer collateral ➔ CA1 synapses

Constitutive α2δ1 deletion alters excitatory synapse number, and α2δ1 overexpression can result in increased mEPSC frequency in dissociated cells ^83^. To disentangle presynaptic α2δ1 roles at CA1 synapses, we bred α2δ1^f/f^ mice with GluK4-cre mice, which express Cre recombinase in CA3 ^55^, generating α2δ1^ΔCA3^ mice (Fig. 2A). As expected, α2δ1 is absent from putative PNs, but preserved in scattered, putative interneurons, in field CA3 in α2δ1^ΔCA3^ mice (Fig. 2B). Importantly, α2δ1 is intact in CA1 PNs (Fig. 2B), confirming that α2δ1^ΔCA3^ mice allow the specific interrogation of presynaptic α2δ1 roles in Schaffer collateral ➔ CA1 PNs synapses, without altering postsynaptic α2δ1 expression (Fig. 2A). Confocal analysis showed no statistical differences in postsynaptic organization in the CA1 SR of α2δ1^ΔCA3^ mice and α2δ1^f/f^ counterparts (Fig 2C, Suppl. Fig. 3). Consistent with this finding, Structured Illumination Microscopy (SIM) imaging showed that the density of excitatory presynaptic (labeled with VGLUT1, Fig. 2D, E) and postsynaptic (labeled with PSD-95, Fig. 2D, F) elements were not altered in CA1 SR of α2δ1^ΔCA3^ mice compared to α2δ1^f/f^ counterparts. Their colocalization was not altered either (Fig. 2D, G). The density of GluA1 puncta (Fig. 2D, H), and the colocalization of GluA1 with VGLUT1 (Fig. 2D, I) and PSD-95 (Fig. 2D, J), was also not altered. Together, these data indicate that presynaptic loss of α2δ1 does not affect gross excitatory synapse organization ^84^, nor does it cause severe mislocalization of AMPARs.

**Figure 2.**
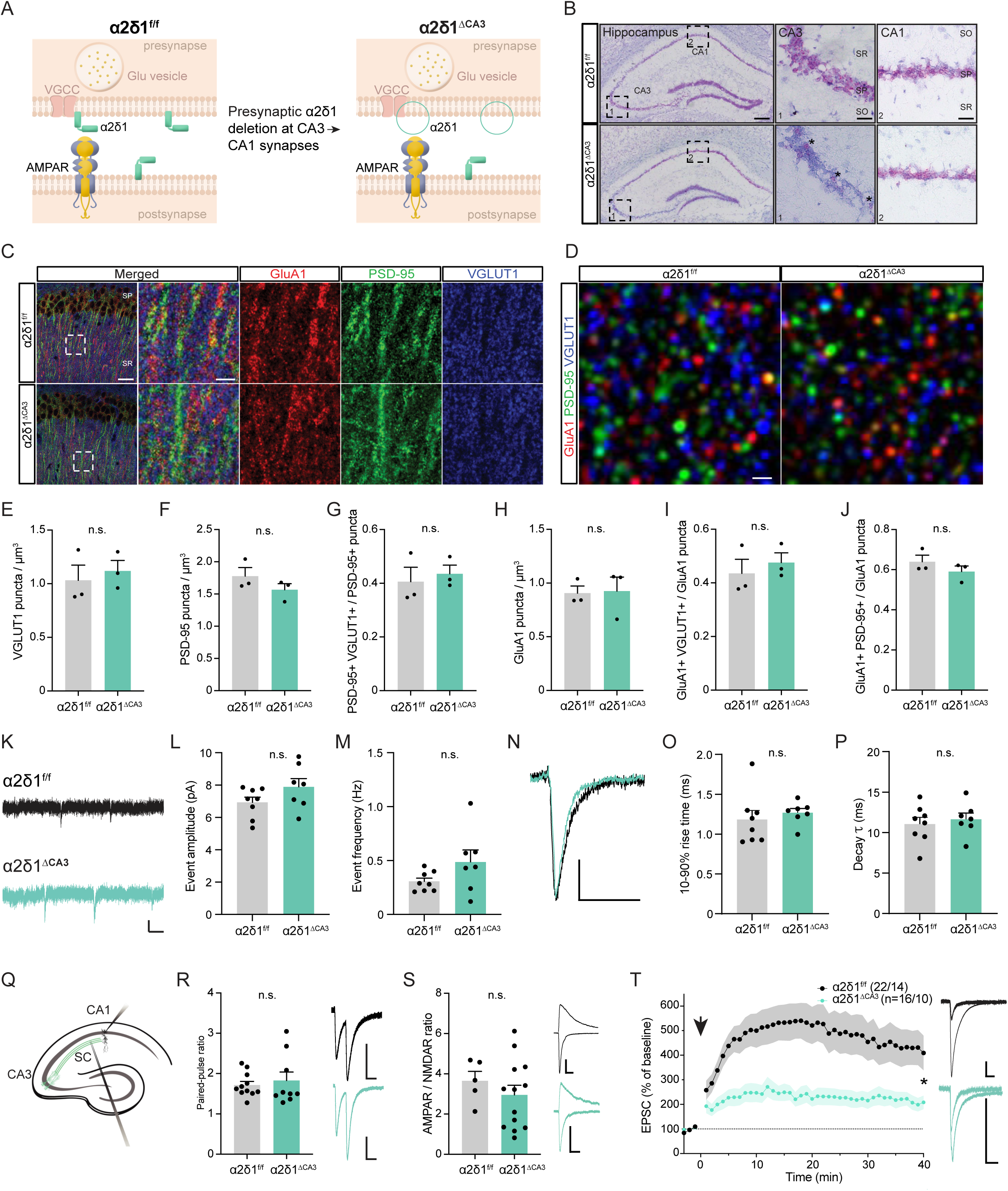
Presynaptic deletion of α2δ1 affects CA3➔CA1 synaptic plasticity. A, Schematic of the presynaptic α2δ1 deletion at CA3➔CA1 synapses, comparing α2δ1^f/f^ (left) with α2δ1^ΔCA3^ (right) mice. B, α2δ1 mRNA ISH in the hippocampus of α2δ1^f/f^ (top) and α2δ1^ΔCA3^ (bottom) mice, showing low magnification (left); CA3 (center) and CA1 (right) photomicrographs. Asterisks identify putative interneurons preserving α2δ1 expression in field CA3 in α2δ1^ΔCA3^ mice. C, Representative immunostaining of GluA1 (red) and PSD-95 (green) and VGLUT1 (blue) in hippocampal field CA1 in α2δ1^f/f^ (top) and α2δ1^ΔCA3^ mice (bottom) samples. Scale bar, 25 µm (10 µm insets). D, Representative Structured Illumination Microscopy (SIM) images of GluA1 (red), PSD-95 (green) and VGLUT1 (blue) in hippocampal area CA1 SR in α2δ1^f/f^ (left) and α2δ1^ΔCA3^ (right) samples. Scale bar, 1 µm. E, F, average density of VGLUT1 and PSD-95 positive puncta, respectively, in CA1 SR. G, Proportion of PSD-95 colocalizing with VGLUT1. H, average density of GluA1-positive puncta. I, J, Proportion of GluA1 colocalizing with VGLUT1 and PSD-95, respectively. K, Representative mEPSC traces for α2δ1^f/f^ (top) and α2δ1^ΔCA3^ (bottom) CA1 PNs. L,M, mEPSC amplitude and frequency, respectively, in α2δ1^f/f^ and α2δ1^ΔCA3^ PNs. N, Representative individual mEPSC traces. O, P, mEPSC 10-90% rise time and decay tau, respectively, in α2δ1^f/f^ and α2δ1^ΔCA3^ CA1 PNs. Q, Schematic of the preparation used for evoked EPSC recordings in R-T. R, S, Paired-pulse ratios (PPR) and AMPAR/NMDAR EPSC ratios in α2δ1^f/f^ and α2δ1^ΔCA3^ CA1 PNs, respectively. T, AMPAR EPSC normalized to the mean AMPAR EPSC amplitude before LTP induction (arrow). AMPAR EPSC current traces from α2δ1^f/f^ (black) and α2δ1^ΔCA3^ (teal) neurons shown to the right of R-T. n=3-8 mice/genotype (C-J), n=5-14 cells/genotype (K-T). Scale bars: 5 pA, 200 ms (K), 2 pA, 50 ms (N), 50 pA, 50 ms (R-T). *, p≤0.05; n.s., not statistically significant, unpaired t-test (E-J), Mann-Whitney U test (L-T). SO, stratum oriens; SP, stratum pyramidale; SR, stratum radiatum.

To assess whether presynaptic α2δ1 affects AMPAR transmission, we turned to electrophysiology. Miniature EPSC (mEPSC) recordings showed no significant change in AMPAR mEPSC amplitude or frequency (Fig. 2K-M). AMPAR mEPSC kinetics revealed no alterations in rise time and decay _τ_ in α2δ1^ΔCA3^ CA1 PNs (Fig. 2N-P). We then obtained evoked AMPAR recordings to further assess the functional consequences of presynaptic α2δ1 loss (Fig. 2Q). We found no difference in paired-pulse ratios (PPR, Fig. 2R) between α2δ1^ΔCA3^ and α2δ1^f/f^ counterparts suggesting that glutamate release is not significantly altered after presynaptic α2δ1 deletion. To assess potential effects in baseline AMPAR transmission, we recorded AMPAR/NMDAR EPSCs ratios. No change in the AMPAR/NMDAR ratios was observed (Fig. 2S). These results, consistent with the lack of impact on NMDAR EPSCs of α2δ1 deletion ^84^ suggest that presynaptic α2δ1 deletion does not alter basal synaptic transmission. Finally, we tested whether activity-dependent AMPAR incorporation into CA1 synapses requires presynaptic α2δ1. Interestingly, we observed that the LTP magnitude, as assessed 40 minutes after induction, was reduced in α2δ1^ΔCA3^ mice compared to α2δ1^f/f^ counterparts (Fig. 2T). Together, these findings indicate that presynaptic loss of α2δ1 leaves constitutive synaptic transmission intact, but affects activity-dependent synaptic AMPAR clustering, at CA3➔CA1 synapses.

### Loss of α2δ1 in CA3 specifically affects CA1-dependent memory tasks

GluA1 plays an important role in spatial memory ^85,86^. To test the functional impact of presynaptic loss of α2δ1 in CA3 PNs we measured object location memory (OLM, Fig. 3A), which relies on hippocampal function, particularly on field CA1 ^87,88^, in male and female α2δ1^ΔCA3^ and α2δ1^f/f^ littermates. During the habituation phase, we found an effect of time, but no effect of genotype in locomotion, with no significant interaction between genotype and time (Suppl. Fig. 4A). Total object exploration time and locomotion during training and testing were not significantly different between genotypes (Suppl. Fig. 4B-E). However, α2δ1^ΔCA3^ male and female mice were unable to discriminate the displaced object, in contrast to α2δ1^f/f^ counterparts (Fig. 3B), suggesting that presynaptic α2δ1 deletion leads to long-term spatial memory deficits. In the novel object recognition task (NOR, Fig. 3C), which involves cortical areas in addition to hippocampus ^69^, α2δ1^ΔCA3^ male and female mice perform at levels comparable to α2δ1^f/f^ counterparts, reflecting the targeted genetic manipulation in α2δ1^ΔCA3^ mice (Fig. 3D). Total object exploration time during training and test was not significantly different between genotypes (Suppl. Fig. 4F, G). α2δ1^ΔCA3^ mice traveled significantly more during the training phase, but not during the test phase (Suppl. Fig. 4H, I). Similarly, α2δ1^ΔCA3^ mice performed comparably to their α2δ1^f/f^ counterparts in the forced alternation Y-maze, which also engages multiple brain regions (Fig. 3E-F). Total number of crossings was similar across genotypes (Suppl. Fig. 4J). Overall, these results suggest that α2δ1 deletion in CA3 leads to CA1-specific spatial memory deficits.

**Figure 3.**
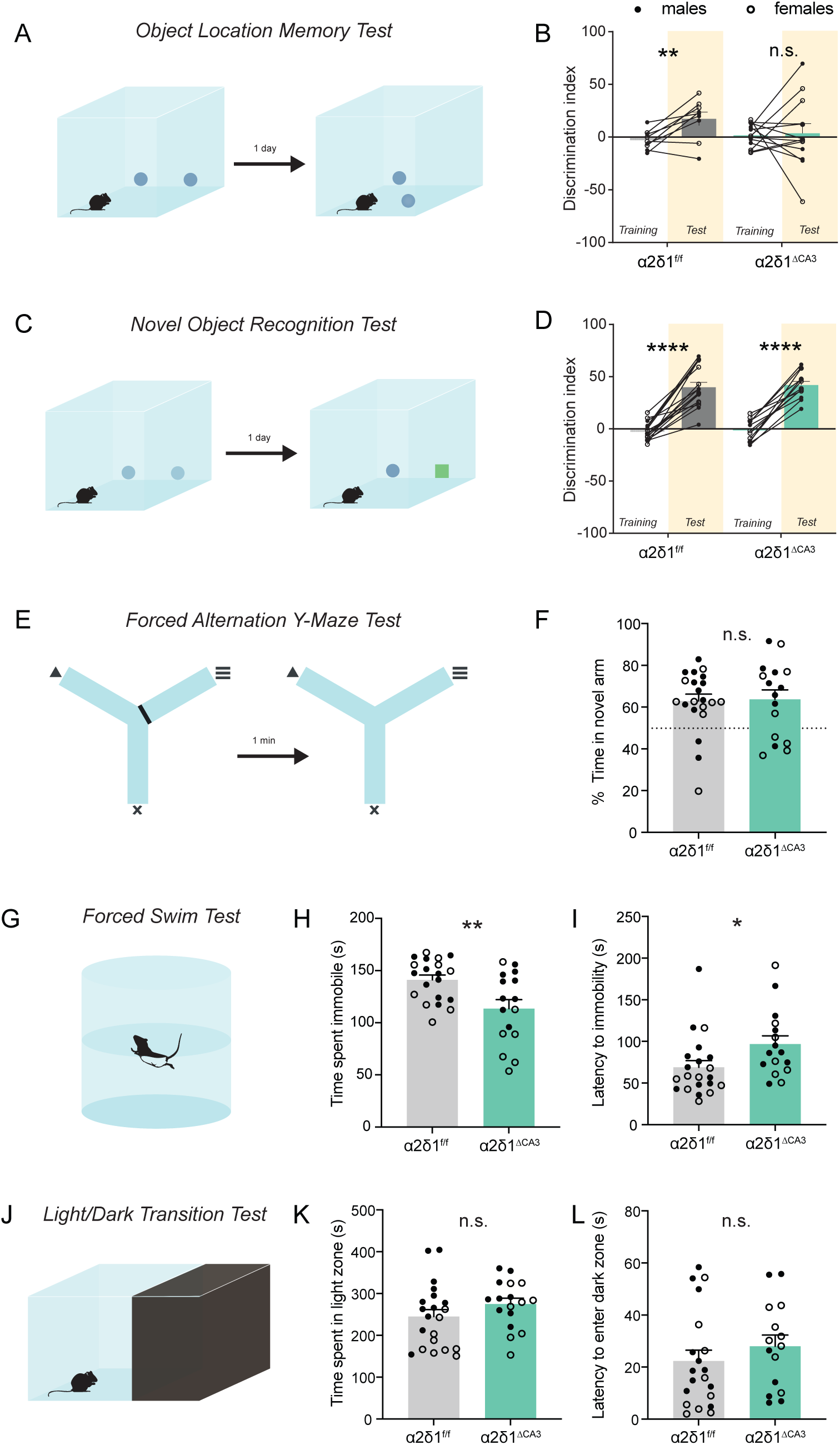
α2δ1^ΔCA3^ mice exhibit deficits in CA1-dependent object location memory and FST. A, Schematic of OLM task. B, Discrimination index in OLM training and test in α2δ1^f/f^ and α2δ1^ΔCA3^ mice. α2δ1^f/f^, α2δ1^ΔCA3^. C, Schematic of NOR task. D, Discrimination index in NOR training and test in α2δ1^f/f^ and α2δ1^ΔCA3^ mice. α2δ1^f/f^, α2δ1^ΔCA3^. E, Schematic of forced alternation Y-maze task. F, Percentage of time spent in novel arm relative to total time spent in novel and familiar arm for α2δ1^f/f^ and α2δ1^ΔCA3^ mice. G, Schematic of forced swim test (FST). H, I, Time spent immobile and latency to immobility, respectively, in α2δ1^f/f^ and α2δ1^ΔCA3^ mice. J, Schematic of light/dark transition test. K, Mean time spent in the light compartment of α2δ1^f/f^ and α2δ1^ΔCA3^ mice. L, Mean latency to enter the dark compartment for α2δ1^f/f^ and α2δ1^ΔCA3^. Filled dots represent male mice; empty circles represent female mice. n=9-21. *, p<0.05; ****, **, p<0.01; ****, p<0.0001; n.s., non-statistically significant, paired t-test (B, D), unpaired t-test (F-L).

### Affective and anxiety-related behaviors are affected in α2δ1^ΔCA3^ mice similarly to GluA1 KO mice

We next tested whether the loss of α2δ1 in CA3 affects other GluA1-dependent behaviors. The forced swim test (FST), which is used to measure learned helplessness in rodents, is altered in GluA1 KO mice ^89,90^. We found that α2δ1^ΔCA3^ mice spent less time immobile in the FST compared to their α2δ1^f/f^ counterparts (Fig. 3G, H). Consistently, latency to immobility was greater in α2δ1^ΔCA3^ mice relative to their α2δ1^f/f^ counterparts (Fig. 3I). Interestingly, α2δ1^ΔCA3^ females displayed a stronger phenotype than males in the FST. α2δ1^ΔCA3^ mice spent comparable time in the light compartment in the light/dark transition test, which is generally used to measure anxiety-like behavior in rodents (Fig. 3J, K). Latency to enter the dark zone in α2δ1^ΔCA3^ mice was comparable to α2δ1^f/f^ littermates (Fig. 3L), and so was the number of entries in light zone (Suppl. Fig. 4K). In this case, analysis of sex differences revealed that α2δ1^ΔCA3^ female mice spent significantly more time in the light compartment than their α2δ1^f/f^ counterparts suggesting, in combination with the FST data, a possible sex-specific effect of α2δ1 function in CA3 in anxiety- and depression-related behaviors. Altogether, these results indicate that behavioral changes in α2δ1^ΔCA3^ recapitulate some of the phenotypes of GluA1 KOs ^91^.

### α2δ1^ΔCA3^ mice show increased latency to PTZ-induced seizures

Our data suggests that SC➔CA1 PN synaptic plasticity is reduced in α2δ1^ΔCA3^ mice. A role for α2δ1 in the pathogenesis of some forms of epilepsy has been proposed, and loss-of-function mutations in the CACNA2D1 gene have been identified in human individuals with epilepsy ^73^. Furthermore, α2δ1 is a molecular target of gabapentinoid drugs (such as gabapentin and pregabalin), that have an antiepileptic profile. All these points suggest that α2δ1 may play an important role in some seizures, but the precise cell types and circuits regulated by α2δ1 presence in the context of epilepsy are unknown. To test the role of α2δ1 in CA3 PNs, we tested whether presynaptic α2δ1 deletion in CA3 PNs affects seizure susceptibility in α2δ1^ΔCA3^ mice. Pentylenetetrazol (PTZ) is a GABA_A_ receptor antagonist commonly used to induce seizures experimentally ^70,92,93^. We administered sub-convulsive (22.5 mg/Kg, IP) doses of PTZ IP until generalized tonic-clonic seizures ensued (Fig. 4A). We observed a significant increase in the cumulative dose necessary to reach tonic-clonic seizures in α2δ1^ΔCA3^ mice compared to α2δ1^f/f^ littermates (Fig. 4B). Similarly, the latency to seizures was increased in α2δ1^ΔCA3^ mice (Fig. 4C). This data suggests that altering transsynaptic AMPAR alignment and subsequent defective synaptic plasticity at SC➔CA1 synapses (and/or CA3➔CA3 recurrents) in α2δ1^ΔCA3^ mice, can reduce hippocampal excitability.

**Figure 4.**
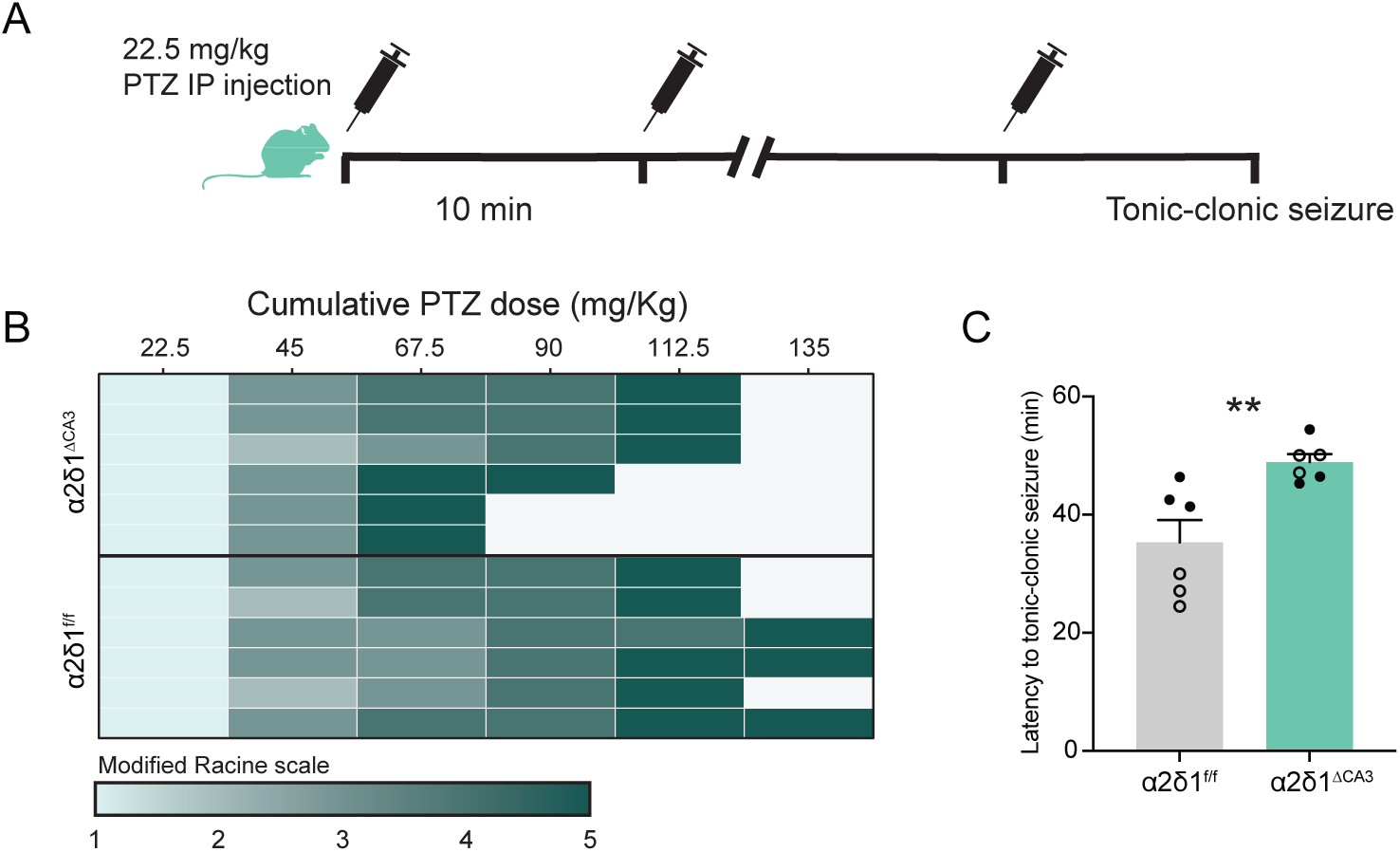
Decreased seizure susceptibility in α2δ1^ΔCA3 mice^. A, PTZ administration timeline. B, Individual Racine stage response and mortality of α2δ1^f/f^ and α2δ1^ΔCA3^ mice following PTZ injections. C, Mean latency to tonic-clonic seizure for α2δ1^f/f^ and α2δ1^ΔCA3^ mice. Filled dots represent male mice; empty circles represent female mice. n=6 per genotype. **p≤0.01, unpaired t-test.

## Discussion

AMPARs have relatively low affinity for glutamate. Therefore, AMPAR EPSC size is determined by the concentration of AMPARs within ∼100 nm of glutamate release sites. Consequently, AMPAR nanoclusters are predominantly observed in alignment with presynaptic glutamate release sites ^12,13,15,17,18,94,95^. However, the mechanisms underlying transsynaptic AMPAR organization are not well understood yet.

Here we identified the extracellular VGCC auxiliary subunit α2δ1 as a GluA1 ATD interacting protein. Molecular docking simulations with α2δ1 in association with VGCC α and β subunits suggest that transsynaptic α2δ1/GluA1 ATD interactions can occur, presumably tuning transsynaptic AMPAR alignment with release sites. Supporting this possibility, presynaptic loss of α2δ1 affects postsynaptic AMPAR-dependent synaptic plasticity, excitability and spatial memory at CA3➔CA1 synapses. To note, α2δ1 can also localize independently of VGCC ^84,96,97^, and our data indicates that the GluA1 ATD can also bind free α2δ1. Interestingly, other proteins initially identified as VGCC subunits, notably stargazin (TARP γ-2), were later found to play a prominent role as AMPAR auxiliary proteins ^11^. In any case, our findings expand the list of calcium channel-independent actions of α2δ subunits ^81,96^, which include regulating synaptogenesis ^98,99^ and acting as synapse organizers ^15,77,78,97,100^.

ATD truncation does not affect AMPAR function ^37^, but has subunit-specific effects on synaptic localization: ATD-lacking GluA1 subunits, but not GluA2 subunits, distribute more diffusely than full length subunits in the postsynaptic membrane ^9^, and activity-dependent synaptic docking of GluA1, but not GluA2, relies on its ATD, at least at SC➔CA1 PN synapses ^37,38,40^. On the other hand, GluA2 ATDs, which occupy central positions in the channel, contribute to stabilizing the AMPAR ATD structure and may facilitate constitutive synaptic AMPAR incorporation ^48^. Interestingly, α2δ1 enables synaptic docking of AMPARs recruited during activity-dependent processes but does not influence constitutive AMPAR-mediated transmission. These data are consistent with a two-step model for AMPAR stabilization at synapses: initially, transient extracellular α2δ1 / GluA1 ATD interactions facilitate subsynaptic localization of newly recruited AMPARs; subsequent TARP/MAGUK interactions stabilize AMPAR complexes at the optimal locations, making the α2δ1 contribution dispensable.

Synaptic trafficking of GluA1-containing AMPARs underlies LTP at CA1 synapses ^24,25,29^. GluA1 is also required for hippocampus-dependent forms of learning and memory, including spatial working memory, but not for spatial reference memory ^101^. Interestingly, the phenotypes we observed in α2δ1^ΔCA3^ mice, including decreased spatial memory performance and reduced immobility in the FST are consistent with the regulation of GluA1-dependent processes. Similarly, GluA1-containing AMPARs participate in synaptic strengthening during status epilepticus ^102^. Consistently, the GRIA1 gene transcript, which encodes GluA1, is upregulated in human subjects with temporal lobe epilepsy ^103^, and GluA1 KOs have reduced susceptibility to status epilepticus ^104^. Interestingly, CACNA2D1, the gene that encodes α2δ1, is associated with epilepsy and neurodevelopmental disorders ^74,96,105^. Increased latency to PTZ-induced seizures in α2δ1^ΔCA3^ mice is therefore consistent with impaired activity-dependent synaptic AMPAR trafficking in the absence of α2δ1. Gabapentin, pregabalin and other gabapentinoid drugs are known to target α2δ1 ^106,107^. They are widely used for the management of seizures and neuropathic pain. Gabapentin affects activity-dependent AMPAR trafficking, including LTP in corticostriatal synapses ^108^, and may induce, in some cases, cognitive and mood alterations ^109,110^. Our data indicates that mechanisms of gabapentinoids include altering α2δ1’s regulation of activity-dependent synaptic AMPAR clustering.

In summary, these studies provide evidence for a transsynaptic role of auxiliary VGCC channel subunits in activity-dependent AMPAR trafficking, which contributes to cognitive function and circuit excitability. Our work adds to an emerging model in which synaptic AMPAR docking relies on multifarious interactions involving both intracellular as well as extracellular slots. The interrelations between these mechanisms, as well as the activity-dependent regulation of slot availability, constitute exciting topics of future research.

### Limitations of this study

Experimental validation will be required to confirm the transsynaptic α2δ1 / AMPARs interactions predicted by molecular docking experiments.

A possible compensation by other α2δ isoforms has not been explored here. Similarly, whether α2δ1 expression or availability at the presynaptic membrane is activity-regulated, and the underlying mechanisms, remain open questions.

Other extracellular AMPAR interacting proteins identified here, which we did not analyze in detail, may also play a role as extracellular AMPAR slots. Among these, we found the endogenous α2δ1 ligands thrombospondins, among the extracellular GluA1 ATD interactors in our proteomics assay, providing additional support for the involvement of this signaling pathway. Other proteins include extracellular matrix glycoproteins which have been shown to be involved in AMPAR trafficking and synaptic plasticity ^111^.

Finally, although in this study we characterized α2δ1 function in GluK4-cre expressing cells, mainly CA3 PNs, its function in other cells and brain regions remains to be determined.

## Acknowledgments

We would like to thank Dr. Roger Nicoll for supporting initial experiments of this project in his laboratory. Mulatwa T. Haile, Dr. Johanna Extrémet, Alexis Madrid, Michael Nguyen, and Dean Madali for their help with experiments and the Diaz-Alonso lab members for fruitful discussions. Dr. Christine Gall for extensive comments on this manuscript. Dr. Jessica Childs and Dr. Marcelo Wood for advice and support with Basescope-ISH experiments. This work was supported by grants K99/R00 MH118425 and BBRF 30264 and UCI start-up funds (J.D.-A.). G.S. is supported a the T32 Training Program in Epilepsy Research (T32NS045540). M.A.S. is supported by a Eugene Cota-Robles fellowship and the Howard Schneiderman T32 Training Program in Learning and Memory (#T32MH119049). The UCSF proteomics core is supported by the Dr. Miriam and Sheldon G. Adelson Medical Research Foundation (KL & ALB). The Optical Biology Core Facility of the Developmental Biology Center is supported by grants CA-62203 and GM-076516.

## Author contributions

G.S. performed and analyzed electrophysiology experiments; K.L. performed MS experiments and A.B., J.J. R-F. and J.D.A. analyzed MS results; G. S., E. S. and J. D.-A. performed biochemistry experiments; J.J. R-F. performed molecular docking simulations; G.S. and M.A.S. performed and analyzed histology experiments; G.S. and A.V.K. performed and analyzed behavior experiments, with equipment and guidance from L.Y.C; G.S and J.D.-A. drafted, and all authors edited the manuscript. J.D.-A. conceptualized and coordinated the study.

## Supplementary Figure legends

**Suppl. Fig. 1. Analysis of proteomics data.** Flow of the analysis of unbiased GluA1 ATD and GluA2 ATD interacting proteins. See Suppl. Table 1 for full list, specific and unique GluA1 ATD and GluA2 ATD interacting proteins.

**Suppl. Fig. 2. Alignment of α2δ1 sequences in different species in the putative GluA1 ATD contact surface.** A, Alignment of human (purple), rabbit (green), mouse (red) and rat (yellow) α2δ1 sequences involved in a putative interaction with GluA1 ATD. PDB or Uniprot IDs are shown. B, Chimerax superimposition of α2δ1 models from different species shows that the GluA1 ATD interaction interface is largely conserved. Color coding is as in A: purple, 7vfv (human); dark blue, 7vfv (human, after docking); green, 6jp8 (rabbit); red, mouse (AF3 from Uniprot O08532), yellow, rat (AF3 from Uniprot P54290). Light blue is GluA1 and light tan is GluA2 (6njl, rat). C, Top and side views of B, highlighting some of the residues involved in the putative interaction with the GluA1 ATD (labelled as residue# based on 7vfv). Color coding as in B.

**Suppl. Fig. 3. Confocal analysis of excitatory synapse density in CA1 SR in α2δ1^f/f^ and α2δ1^ΔCA^**^3^ **mice.** A, average density of GluA1 positive puncta. B, average density of PSD-95 positive puncta. C, proportion of GluA1 puncta colocalizing with PSD-95. n=7-8 mice/genotype. n.s., non-statistically significant, unpaired t-test.

**Suppl. Fig. 4. Control behavioral assessments in α2δ1^f/f^ and α2δ1^ΔCA^**^3^ **mice.** A, Mean distance traveled during habituation phase for α2δ1^f/f^ and α2δ1^ΔCA3^ mice. B, C, Mean object exploration time during training (B) and test (C) for α2δ1^f/f^ and α2δ1^ΔCA3^ mice in OLM task. D, E, Mean distance traveled during training (D) and test (E) for α2δ1^f/f^ and α2δ1^ΔCA3^ mice in OLM task. F, G, Mean object exploration time during training (F) and test (G) for α2δ1^f/f^ and α2δ1^ΔCA3^ mice in NOR task. H, I, Mean distance traveled during training (H) and test (I) for α2δ1^f/f^ and α2δ1^ΔCA3^ mice in NOR task. J, Total number of arm entries for α2δ1^f/f^ and α2δ1^ΔCA3^ mice in the forced alternation Y-maze task. K, Total number of entries into the light zone for α2δ1^f/f^ and α2δ1^ΔCA3^ mice in the light-dark box. n=9-21 mice/genotype. **p≤0.01; n.s., non-statistically significant, unpaired t-test.

## Supplementary Table legends

**Suppl. Table 1. Specific and unique GluA1 ATD and GluA2 ATD binding proteins identified with mass spectrometry.** Table shows specific (present in GluA1 or GluA2 ATD beads but absent in control beads) and unique (present in GluA1-specific but absent in GluA2-specific or vice-versa) GluA1 ATD and GluA2 ATD interacting proteins in synaptosomal-enriched P2 fractions from mouse brain identified with mass spectrometry. Protein identifier, name, sumed normalized spectral abundance factor (NSAF) and Uniprot predicted localization are indicated. See Suppl. Figure 1 for detailed workflow.

**Suppl. Table 2. Energy distribution of residues involved in putative contact surfaces between α2δ1 and GluA1 ATD.** List of residues involved in putative α2δ1 / GluA1 ATD interaction with binding energies. Residue ID, Van der Waals (VdW), electrostatic (ELE), generalized Born (GB) and solvent accessible surface area (SA) energies are indicated. Residues with highest contribution are highlighted in Fig. 1H.

