## Supplementary figures and images for "The VGCC auxiliary subunit α2δ1 is an extracellular GluA1 interactor and regulates LTP, spatial memory, and seizure susceptibility"

### Supplemental Figure 1

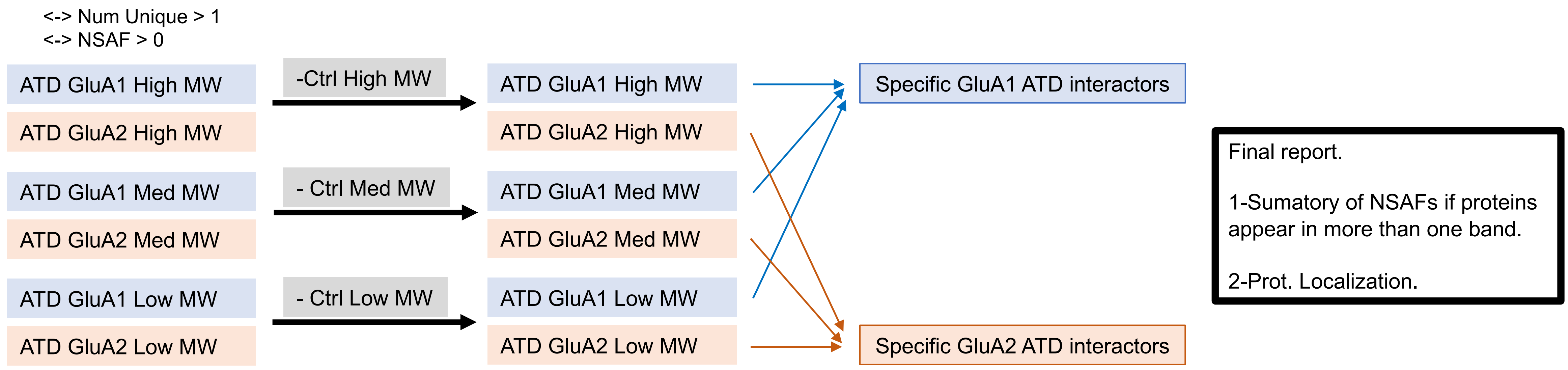

### Supplemental Figure 2

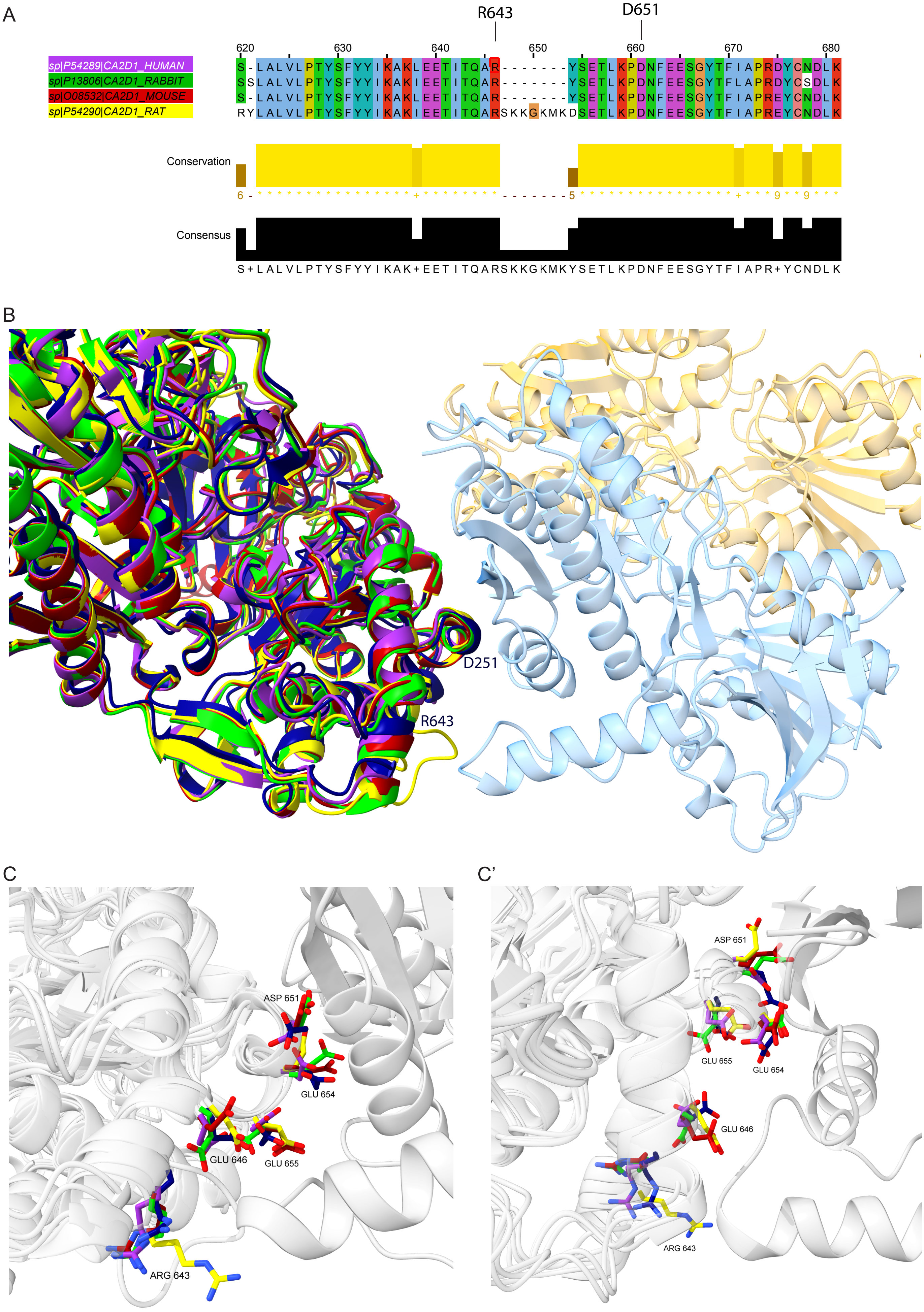

### Supplemental Figure 3

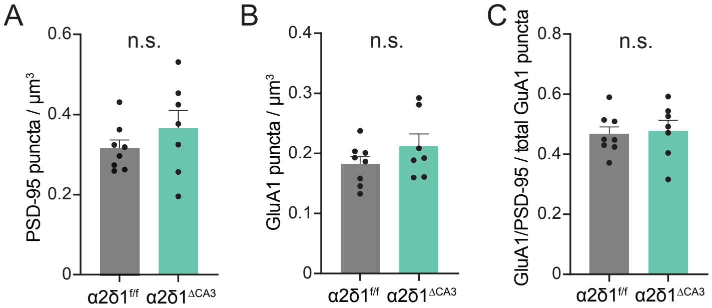

### Supplemental Figure 4

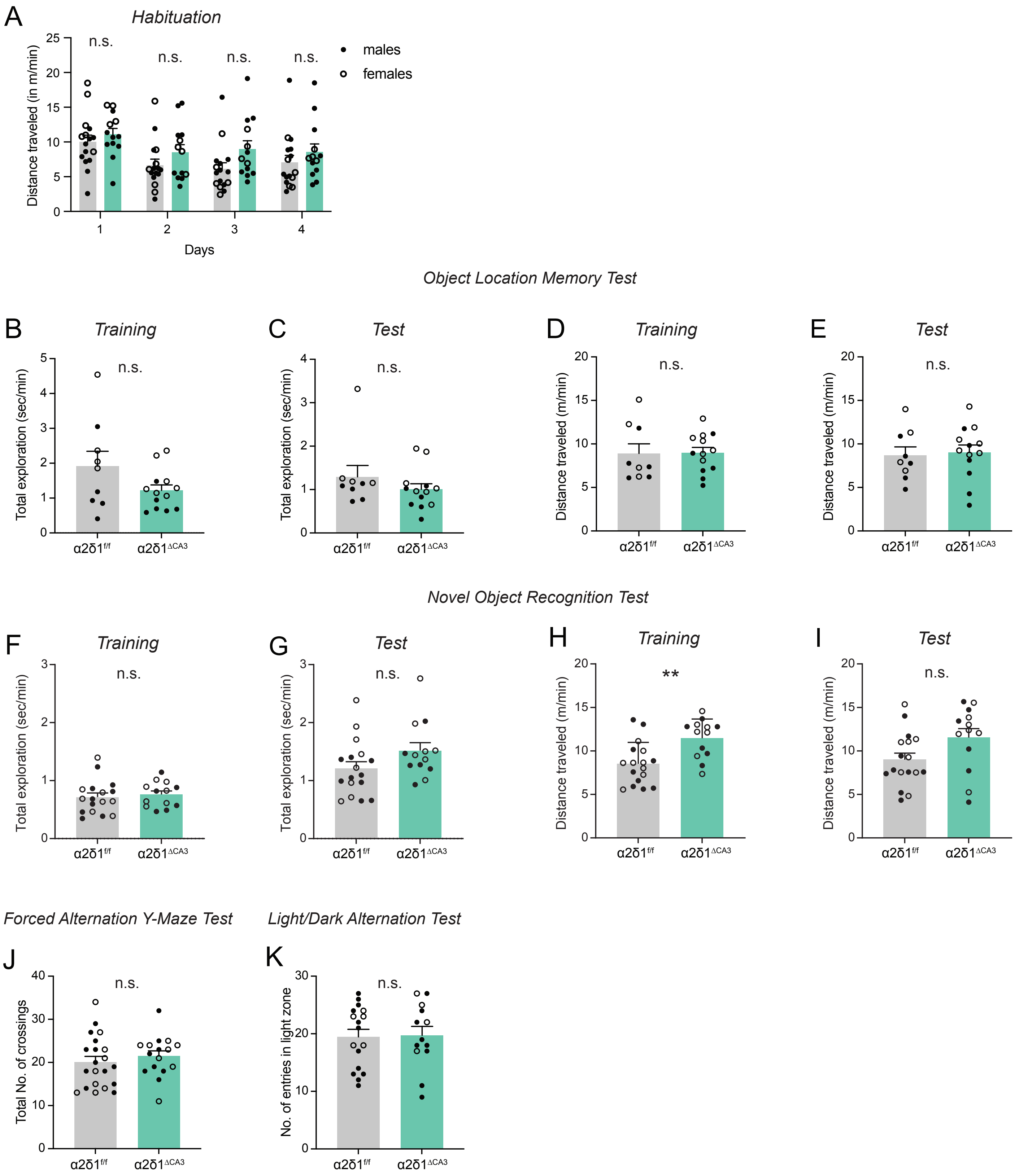
